# Attention periodically samples competing stimuli during binocular rivalry

**DOI:** 10.1101/253740

**Authors:** Matthew James Davidson, David Alais, Naotsugu Tsuchiya, Jeroen J.A. van Boxtel

## Abstract

The attentional sampling hypothesis suggests that attention rhythmically enhances sensory processing when attending to a single (~8 Hz), or multiple (~4 Hz) objects. Here we investigated using binocular rivalry whether attention samples sensory representations that are not part of the conscious percept, during competition for perceptual dominance. When crossmodally cued toward a conscious image, subsequent changes in consciousness occurred at ~8 Hz, consistent with rates of undivided attentional sampling. However, when attention was cued toward the suppressed image, changes in consciousness slowed to ~3.5 Hz, indicating the division of attention away from the conscious visual image. In the electroencephalogram, we found that at 3.5 and 8 Hz, the strength of inter-trial phase coherence over fronto-temporal and parieto-occipital regions correlated with behavioral measures of changes in perception. When cues were not task-relevant, these effects disappeared, confirming that perceptual changes were dependent upon the allocation of attention, and that attention can flexibly sample away from a conscious image in a task-dependent manner.

## Introduction

Recent behavioral and electrophysiological evidence suggests that despite our seamless visual experience, incoming visual information is periodically enhanced for analysis in the visual system (Rufin VanRullen, 2016a, 2016b; Zoefel & VanRullen, 2017). This periodic sampling mechanism is proposed to result from the allocation of visual attention (Busch & VanRullen, 2010; VanRullen, Carlson, & Cavanagh, 2007; Zoefel & VanRullen, 2017), wherein alternating windows of high and low attentional resources operate to parcel incoming visual information, similar to the sequential frames that capture film within a video camera (Chakravarthi & VanRullen, 2012; VanRullen & Dubois, 2011). Whether stimuli are presented at the appropriate phase (Busch, Dubois, & VanRullen, 2009; Mathewson, Gratton, Fabiani, Beck, & Ro, 2009; VanRullen et al., 2007) or location (Dugué, McLelland, Lajous, & VanRullen, 2015; Dugué, Xue, & Carrasco, 2017; Dugué & VanRullen, 2014; Huang, Chen, & Luo, 2015; Landau & Fries, 2012) of this sampling mechanism has been shown to modulate the accurate detection of a visual stimulus, in stark contrast to our experience of an uninterrupted visual environment.

To date, primary neural evidence for the rhythmic gating of visual processing stems from the dependence of target detection on the pre-target phase of neural oscillations at approximately 7-8 Hz (Busch & VanRullen, 2010; Busch et al., 2009). These spontaneous fluctuations in detection may result from the allocation of visual attention toward a single location (Busch & VanRullen, 2010; Dugué et al., 2015; Spaak, de Lange, & Jensen, 2014; VanRullen, 2016a; Zoefel & VanRullen, 2017), and support the assumption that neural excitability cycles gate and filter incoming information for further processing (Schroeder & Lakatos, 2009; VanRullen, 2013; Zoefel & VanRullen, 2017).

This periodic gating of visual perception is also prominent behaviorally in the time-course of detection accuracy. Spectral analyses applied to high temporal resolution behavioral measures reveal 7-8 Hz modulations in performance following cues to reorient attention (Dugué et al., 2015; Fiebelkorn, Saalmann, & Kastner, 2013), which slow proportionately when attention is divided between two or more locations (e.g. Chen, Wang, Wang, Tang, & Zhang, 2017; Holcombe & Chen, 2013; Huang et al., 2015; Landau & Fries, 2012; Landau, Schreyer, van Pelt, & Fries, 2015; VanRullen, 2013). For example, Landau and Fries (2012) observed that following a cue to reorient attention to either the left or right visual hemifield, target detection oscillated at a 4 Hz counterphase rhythm depending on whether cues were congruent or incongruent with the target location. Critically, this counterphase sampling of visual information persisted at ~ 4 Hz when attention was directed to two locations on a single object (Fiebelkorn et al., 2013), and when cues to reorient attention were incongruent with target location – requiring a subsequent shift in the allocation of attention to a second location (Huang et al., 2015). These successive fluctuations in target detection and counterphase sampling between locations have led to the suggestion that an intrinsic ~7-8 Hz attentional rhythm can be allocated over space and time in a sequential manner (Fiebelkorn et al., 2013; A. O. Holcombe & Chen, 2013; Landau & Fries, 2012; VanRullen, 2013; Zoefel & VanRullen, 2017).

Here, we tested if rhythmic attentional sampling is at play during binocular rivalry. During binocular rivalry, incompatible images are presented to each eye which results in stochastic perceptual alternations, with one image visible at a time while the other is suppressed (Alais, 2012; Alais & Blake, 2005; Maier, Panagiotaropoulos, Tsuchiya, & Keliris, 2012). In an experiment designed to induce or delay these transitions using crossmodal cues, we found that changes in consciousness were occurring rhythmically after the reorientation of attention. These fluctuations occurred depending on whether the crossmodal cue directed attention toward either the dominant or suppressed visual image, resulting in ~8 Hz and ~ 3.5 Hz oscillations, respectively. Critically, these rhythms were observed in both behavior and the electroencephalogram (EEG), and were absent when cues were not task-relevant. This approximate halving of frequency suggests that when non-visual input is inconsistent with the ongoing visual percept, attentional sampling can flexibly orient away from a consciously perceived image, seemingly ‘searching for’ alternative sensory information to resolve the conflict.

## Results

### Attending to low-frequency crossmodal stimulation promotes the perceptual dominance of low-frequency flicker during binocular rivalry

We manipulated the conscious visibility of images across two sessions of 24 x 3-minute binocular rivalry blocks. Subjects (*N*=34) continuously reported the content of their visual consciousness via button press to indicate which image they currently perceived, while neural activity was simultaneously recorded via 64-channel EEG (see Methods). Rivalry stimuli were orthogonal sinusoidal gratings which underwent contrast modulation, one at 4.5 Hz and the other at 20 Hz (Figure 1a). In each 3-minute block, we intermittently presented 12 crossmodal cues (mean duration 2.6 s), which were amplitude-modulated signals presented in the auditory and/or tactile modality at a frequency congruent with one of the visual stimuli (4.5 or 20 Hz). Three null cues (visual-only periods) without any crossmodal stimulation were also presented to increase the uncertainty of stimulus timing. The visual-only periods also served as a baseline to compare the behavioral effects of crossmodal cues (see below).

**Figure 1.**
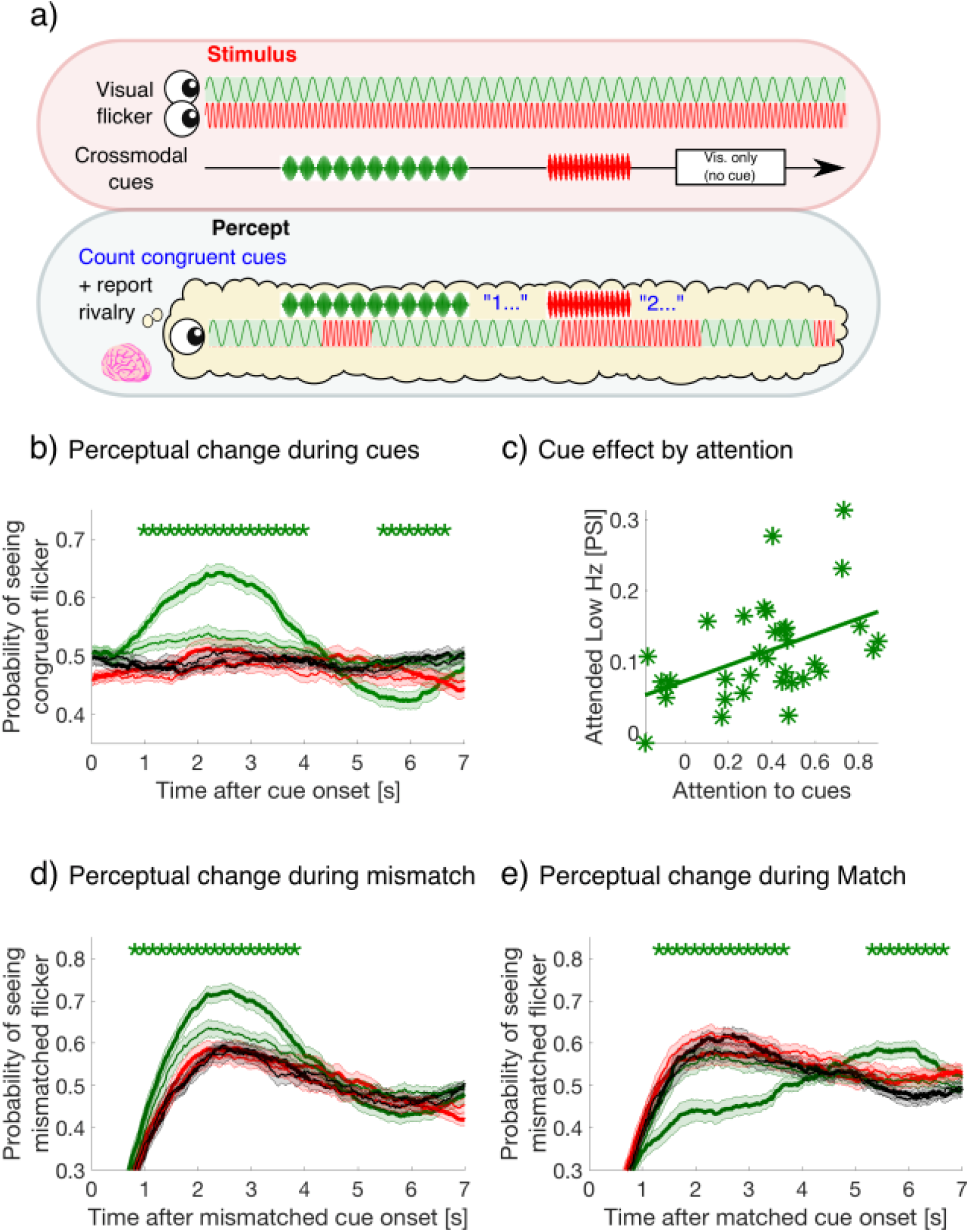
Experimental paradigm and behavioral results. a) A schematic time course showing stimulus presentation and reported visual percept. Each eye was presented with a 4.5 or 20 Hz sinusoidal flicker throughout 3-min blocks. Subjects reported their perceptual state through button press. Crossmodal cues (also 4.5 or 20 Hz; 2, 3.1 or 4s in duration) or visual-only periods (2.6s in duration) were separated by inter-stimulus intervals of 7-10 s. b) Button-press data, aligned at cue onset, were averaged over all crossmodal cue and visual-only periods per subject, then averaged over subjects for each cue condition. Y-axis represents the proportion of button-presses reporting congruent crossmodal and visual flicker at each time point, sampled at 60 Hz (or every 16.7 ms). Colored lines and their shading show mean **±**1 standard error across 34 subjects during attended and ignored cues (thick and thin lines) and low and high frequency (green and red colors). Black lines represent the equivalent probability for visual-only periods, **serving as baseline (Methods).** Asterisks indicate a significant difference between cues at each time point (repeated-measures ANOVA followed by planned comparisons). We use FDR q = .05 for the statistical threshold unless noted otherwise. c) Crossmodal effects are mediated by task-relevant attention. Our measure of crossmodal effects, the perceptual switch index (PSI, y-axis), is defined as the mean difference for the probability of seeing congruent flicker during 1-4 sec after the cue onset for attended-low-frequency cues (solid green in panel b) compared to other cue types. Attention-task performance (x-axis) is the correlation coefficient between the reported and actual congruent stimuli when comparing between rivalry percepts and crossmodal cues at offset (See Methods for details). The across-subject correlation between the two variables was strong (r(32) = .46, p = .006, two-tailed), demonstrating the crossmodal effects were strongly dependent on performance during the attention task. d) and e) Button-press data aligned at cue onset, with lines and shading as in panel b). Y-axis showing the proportion of button-presses reporting the mismatched flicker at each time point, after d) visual-crossmodal mismatch, or e) visual-crossmodal match at cue onset. Only the data of the attended-low-frequency condition differed significantly from the other conditions, including visual only periods.

All cue periods were separated by jittering the ISI between 7-10 s. As a result, the timing of the crossmodal cues were controlled completely independently from the perceptual reports, and were presented at any point relative to the onset of the currently dominant percept (i.e., no closed-loop control).

In order to investigate whether the allocation of attention to crossmodal cues alters the contents of visual consciousness during binocular rivalry, we varied attentional instructions over two sessions of the experiment. For one of their two sessions (day 1 for *n*=16, day 2 for *n*=18), we asked subjects to count the number of times that the temporal frequency of crossmodal cues coincided with their conscious visual percept at crossmodal cue offset (see Methods). For their other session, subjects were instructed to focus on reporting their visual percept alone – ignoring any crossmodal cues.

Following the onset of a crossmodal cue, the probability of perceiving a congruent visual image increased only during attended low-frequency cues compared to all other cue types, during the period 0.68 to 3.97 s after cue onset (repeated measures ANOVAs followed by planned comparisons, FDR q = .05, Figure 1b). To confirm that this effect was due to attention, we performed a correlation-based behavioral analysis. First, we computed the correlation coefficient (x-axis in Figure 1c), between each subject’s verbally reported number of congruent cues (i.e., their attentional task during attend conditions), to the actual number of cues that were congruent with their visual percepts based on button-press data. Second, we defined the strength of the crossmodal cueing effect for attended low-frequency cues compared to other cue types (y-axis in Figure 1c), as the difference in the probability of seeing the congruent visual flicker during 1 to 4 s after cue onset. We call this the perceptual switch index (PSI), as it reflects the degree of perceptual switch after cue onset. The magnitude of these two variables displayed a strong positive correlation (*r(*32) = .46, *p* = .006, two-tailed), suggesting that the cross-modal cueing effect was indeed mediated by attention.

Due to the ongoing dynamics of binocular rivalry, this cueing effect can be calculated when visual and crossmodal information mismatched or matched at cue onset. When crossmodal cues mismatched with the visual percept at cue onset, the likelihood of switching to the previously suppressed, yet matched visual stimuli significantly increased for attended low-frequency cues compared to all other cue types over a time period from 0.62 to 4.12 s (FDR q = .05, Figure 1d). By contrast, when visual and crossmodal cues matched at cue onset, the effect of attending to low-frequency crossmodal cues delayed changes to the previously suppressed visual percept compared to all other cue types, over the period from 1.05 to 3.58 s (FDR q = .05, Figure 1e). Comparison against the visual-only cue period yielded the same conclusion, confirming that the attended low-frequency cues significantly influenced rivalry dynamics, while other cue types did not. As the overall crossmodal effects were unique to the attended low-frequency condition, we focused our subsequent attentional sampling and EEG analysis on this condition.

### Binocular rivalry dynamics after mismatched and matched crossmodal cues

Our previous analysis showed that relative to matched crossmodal cues, mismatched crossmodal cues lead to more perceptual switches, as the visually perceived image changed to become congruent with the crossmodal input. In the context of the attentional sampling hypothesis, we directly tested if these changes were occurring rhythmically after the reorientation of attention, and specifically investigated the timing of the *first* switch after cue onset, defined as the first change in button-state after cue onset.

To determine if cues affected the timing of first switches, we calculated the cumulative density function of each subject’s first switches after cue onset (Figure 2a). Compared to visual-only cue periods, first-switches after cue onset occurred earlier for mismatched cues, indicating an earlier change to the congruent, previously suppressed, visual flicker. By contrast, following matched cues first-switches during rivalry were delayed, indicating an extended maintenance of the congruent visual percept when matched with attended low-frequency crossmodal cues. The facilitation of switches by mismatched cues was observed from 0.63 to 2.45 s and 3.78 to 6.87 s relative to cue onset, with matched cues delaying switches from 1.27 to 3.77 s after onset (paired samples *t*-tests, FDR q = .05, in Figure 2b).

**Figure 2.**
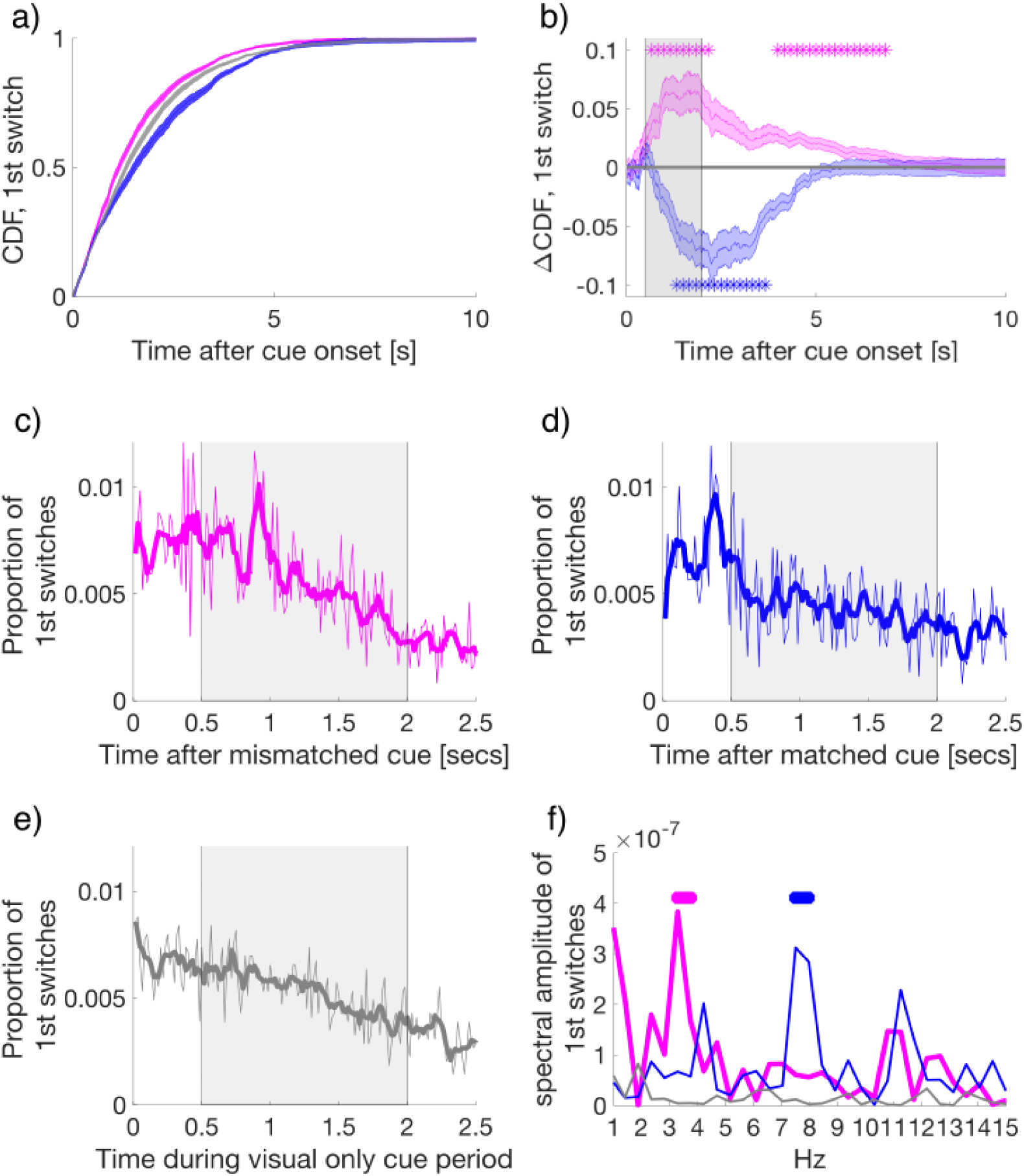
a) The cumulative density function (CDF) of the time to first-switch. Mismatched, matched, and visual-only conditions are colored in magenta, blue, and grey in all panels a-f. Lines and shading show mean and standard error across subjects (N=34) for a and b. b) The difference in CDFs between conditions. Asterisks mark statistical significance (paired-samples t-tests) comparing mismatched or matched cues to visual-only periods. FDR q = .05. c-e) The time course of the proportion of first switches made after cue onset in c) mismatched, d) matched, and e) visual-only conditions. Thin lines show the mean proportion of first-switches, binned in 16.7 ms increments and averaged across subjects. Thick lines show the smoothed data for visualization. Grey shaded regions show the time window used for spectral analysis in f). f) The amplitude spectra for the time course of switches in conditions in c-e). Asterisks indicate significant clusters (at least two neighboring frequency bins) after permutation and cluster-based corrections for multiple comparisons (see Methods). The permuted null distribution and critical value for the identified clusters in f) are shown in Figure S3)

After cue onset, the time-course for the probability of first switches displayed oscillatory patterns for mismatched and matched conditions (Figure 2c and d), but not the visual only condition (Figure 2e). Each data point represents the proportion of first switches which occurred at each time bin (16.7 ms intervals), calculated first per individual, and then averaged across subjects (subject-level data is shown in Figure S1).

To quantify these patterns, we applied the Fourier transform to the period 0.5 to 2 s after cue onset (skipping the first 0.5 s to avoid an onset transient, see Figure S2) as performed by previous investigations of attentional sampling (Laura Dugué et al., 2015; Laura Dugué, Roberts, & Carrasco, 2016; Fiebelkorn et al., 2013; Landau & Fries, 2012). For this analyses, we corrected for multiple comparisons by using non-parametric cluster-based permutations (Maris & Oostenveld, 2007), with thresholds set to *p* < .005 (Benjamin et al., 2017) for identification within a cluster, and a final critical value for significance set to *p* = .05, cluster corrected (see Methods).

Strikingly, when the temporal frequency of the cue matched the conscious visual flicker at cue onset, the first perceptual switches followed a 7.5-8 Hz rhythm (*p_cluster_* < .001, Figure 2f blue), consistent with findings that show attention samples from single locations at a rate of approximately 7-8 Hz (Laura Dugué et al., 2015; Fiebelkorn et al., 2013). However, when crossmodal cues were mismatched with the dominant visual image at cue onset, the amplitude spectrum of perceptual switches peaked between 3.3-3.75 Hz (*p_cluster_* < .001, Figure 2f magenta). This slower rhythm of perceptual changes is consistent with findings that show attention samples two locations at a rate of approximately 3.5-4 Hz (Fiebelkorn et al., 2013; Landau & Fries, 2012; Landau et al., 2015). No significant peaks were detected for the visual only condition (Figure 2f, gray). As to the remaining three cue combinations (attended high-, ignored low- and ignored high-frequency cues), all failed to exhibit any significant crossmodal effects on perceptual switches compared to visual only periods (shown Figure 1b, d, e, and Figure S4). Thus, we did not pursue further spectral or neural analyses of these conditions.

### The neural correlates of divided and focused attentional sampling

We hypothesized that at our behaviorally observed attentional sampling frequencies (3.5 and 8 Hz), we should be able to identify the neural correlates of attentional sampling in the EEG signal using an inter-trial phase coherence (ITPC) measure. Previously, the phase of ongoing cortical oscillations have been shown to be reset by external crossmodal events (Frey, Ruhnau, & Weisz, 2015; Lakatos et al., 2009; Mercier et al., 2013; Romei, Gross, & Thut, 2012; van Atteveldt, Murray, Thut, & Schroeder, 2014) and to modulate the probability of target detection (Busch et al., 2009; Landau et al., 2015; Mathewson et al., 2009; Thorne & Debener, 2014; VanRullen et al., 2007). To isolate the specific neural correlates of attentional sampling we compared the evoked ITPC, the increase in ITPC during 0 to 2 s after onset compared to −2 to 0 s before onset, in mismatched and matched cue conditions at the attentional sampling frequencies (3.5 and 8 Hz). Importantly, in these conditions, the physical sensory input was identical (i.e., attending low-frequency tones during binocular rivalry), with the only difference between cues being the subject’s percept at cue onset. Thus, any differences between conditions reflect differences due to crossmodal mismatch or match with the subjective visual percept.

For this analysis, we retained electrodes only after identification of a significant effect (*p* < .05, uncorrected) which also satisfied a spatial cluster-based criterion for selection (Maris & Oostenveld, 2007), and used non-parametric permutation distributions to control for multiple comparisons Maris & Oostenveld, 2007; Figure S5). We found that the mismatched cues induced stronger ITPC than the matched cues, at 3.5 Hz over right fronto-central-temporal electrodes [FT8, C6] (Figure 3a) and at 8 Hz over right parietal-occipital electrodes [P6, PO8] (Figure 3e). Figures 3b and f compare the evoked ITPC spectra in these regions based on mismatched and matched subjective percepts at cue onset, and confirm that our time window was long enough to distinguish the 3.5 from 4.5 Hz stimulus response (with half bandwidth = 0.5 Hz to resolve the frequency of interest).

**Figure 3.**
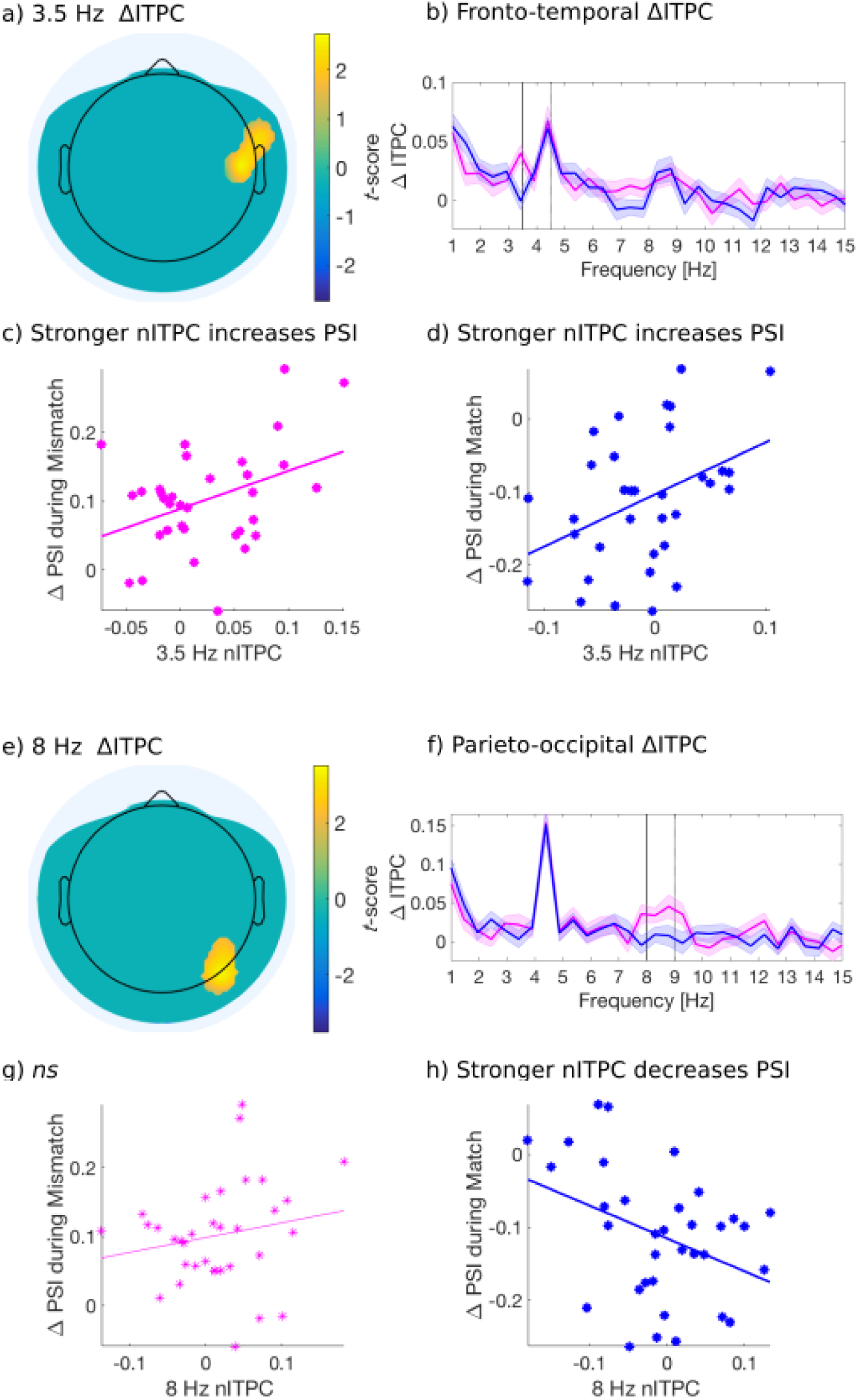
Evoked ITPC at 3.5 Hz (a-d) and 8 Hz (e-h) mediates the probability of switches during rivalry. a, e): Significant differences in evoked ITPC between mismatched and matched cue conditions (multiple comparisons corrected using a cluster-based criterion; Methods). b, f): Evoked ITPC spectra at significant regions in (a) and (e). The magenta and blue lines and their shading show mean ±1 standard error of the mean across 34 subjects for mismatched and matched cues, respectively. Solid vertical black lines mark in (b) the behaviorally observed attentional sampling frequency at 3.5 Hz, and in (f) the 8 Hz sampling frequency observed behaviorally. (c, d): Stronger 3.5 Hz nITPC correlates with increased PSI during (c) mismatched and (d) matched conditions. The x and y-axes represent the normalized ITPC and perceptual switch index respectively (see text for definitions). (g, h): Stronger 8 Hz nITPC correlates with a decreased PSI for (h) matched, but not the (g) mismatched condition.

### Attentional-sampling ITPC strength predicts perceptual outcome

Next, we investigated whether the evoked ITPC at the attentional-sampling frequencies in the above-identified regions (Figures 3a and e) predicted the magnitude of behavioral effects across subjects, shown in Figure 1d-e. We again computed the difference in behavioral effects when comparing attended low-frequency to all other cue types (PSI; 2:4 s after cue onset), as a measure for the degree of perceptual change following mismatched and matched cues. Note that when considering a wider time-window (0:4s for behavioral effects, data not shown) a similar pattern of results was obtained, though weaker due to the lack of differences between cue types in early cue periods (i.e. 0:1s, cf. Figure 1d-e). We used the evoked ITPC from 0 to 2s after cue onset to restrict our analysis to within attended crossmodal cueing periods (which were 2, 3.1 and 4s in duration), and to capture the period where the majority of first switches were made after cue onset (Figure 2c and d). Similar to the PSI, we also subtracted the evoked ITPC across all other conditions from those in the attended low-frequency condition, and abbreviate this as the normalized ITPC (nITPC) below.

In the right fronto-central-temporal electrodes ([FT8, C6]) which significantly differed in 3.5Hz ITPC based on mismatched or matched percepts (Figure 3a), we found that 3.5 Hz nITPC and PSI were positively correlated for both mismatched (*r*(32) = .39, *p* = .024, two-tailed, Figure 3c), and matched cue types (*r*(32) = .40, *p* = .018, two-tailed, Figure 3d). Indicating that for both mismatched and matched cues, increases in 3.5 Hz nITPC facilitated a change in visual consciousness across subjects (Figure 3c-d).

In the parieto-occipital electrodes ([P6, PO8]), we found that 8 Hz ITPC was not correlated with the PSI for mismatched cues (Figure 3g). However, 8 Hz ITPC was negatively correlated with the PSI during matched cues (*r*(32) = -.38, *p* = .026, two-tailed, Figure 3h), demonstrating that increased 8Hz nITPC resulted in fewer perceptual switches across subjects.

## Discussion

Our findings provide novel evidence that attentional sampling exists during binocular rivalry, demonstrated in both behavior and the electroencephalogram (EEG). Behaviorally, we replicated previous evidence that crossmodal cues can cause a switch to previously suppressed visual stimuli when mismatched with the current percept (to bring about crossmodal congruence), as well as increase the maintenance of a dominant visual image if cues matched perception (Figure 1; Lunghi & Alais, 2015; Lunghi, Morrone, & Alais, 2014). Critically, we found distinct attentional sampling frequencies evident in the time-course of first switches during these cues (Figure 2). When crossmodal cues were incongruent in temporal frequency with the dominant visual stimulus, switches in visual consciousness occurred earlier, and within a distinct ~3.5 Hz rhythm. This 3.5 Hz rhythm is consistent with previous reports of divided attentional sampling between two locations (Fiebelkorn et al., 2013; Landau & Fries, 2012; Landau et al., 2015). However, when crossmodal cues were matched in temporal frequency to the dominant visual stimulus, changes in visual consciousness demonstrated an ~8 Hz rhythm, consistent with periodicities in behavioral measures observed when attending to a single visual location (Dugué et al., 2015; Fiebelkorn et al., 2013). In the EEG (Figure 3), distinct correlates of these divided and focused attentional sampling frequencies emerged over fronto-temporal and parieto-occipital sites, respectively, with ITPC strength at these frequencies correlating with the behaviorally reported change in consciousness across subjects.

Traditionally, top-down, voluntary attention has been thought to have limited control over perceptual dynamics during binocular rivalry (Paffen & Alais, 2011; for bottom-up control, including crossmodal stimulation, see Conrad, Bartels, Kleiner, & Noppeney, 2010; Deroy et al., 2016; Guzman-Martinez, Ortega, Grabowecky, Mossbridge, & Suzuki, 2012; Kang & Blake, 2005; Lunghi & Alais, 2013; Lunghi, Binda, & Morrone, 2010; Lunghi et al., 2014; van Ee, van Boxtel, Parker, & Alais, 2009). Our results clearly show additional dependence on the top-down deployment of attention, as without explicit instruction to attend to crossmodal signals, no facilitatory crossmodal effects emerged (see also Jack & Hacker, 2014; Talsma, Senkowski, Soto-Faraco, & Woldorff, 2010; van Ee et al., 2009). This interaction between low-level stimulus features (temporal frequency) and the allocation of attention indicates the facilitative role of both crossmodal stimuli (Deroy et al., 2016; Deroy, Chen, & Spence, 2014) and attention for perceptual transitions during binocular rivalry (Kevin C. Dieter, Brascamp, Tadin, & Blake, 2016; Kevin Conrad Dieter & Tadin, 2011; Paffen & Alais, 2011; P. Zhang, Jamison, Engel, He, & He, 2011), here revealing a previously unknown periodicity to changes in visual perception.

Previous investigations of attentional sampling have relied upon a brief cue to reorient attention, before estimating the time-course of target detection by densely sampling subject responses over closely spaced target-presentation intervals. Our design is unique in that ‘target-detection’ here is operationalized as the first reported change in visual consciousness for a continuously presented stimulus, resolved at 16.7 ms (or 60 Hz) from 500 ms to 2000 ms following cue-onset.

Past research has demonstrated approximately 7-8 Hz fluctuations in perceptual performance following the allocation of visual attention to a single location (Laura Dugué et al., 2015; Fiebelkorn et al., 2013; R. VanRullen et al., 2007; Zoefel & VanRullen, 2017), commensurate with suggestions that cortical oscillations at approximately 7-8 Hz gate the content of visual perception (Busch & VanRullen, 2010; Hanslmayr, Volberg, Wimber, Dalal, & Greenlee, 2013). In our binocular rivalry paradigm, we also observed changes in visual consciousness occurring within an 8 Hz rhythm, yet unique to when cues were congruent with the dominant visual stimulus at cue onset. By contrast, perceptual sampling has previously been observed at ~4 Hz when cues have encouraged dividing attention between two objects or locations (Fiebelkorn et al., 2013; Huang et al., 2015; Landau & Fries, 2012; Landau et al., 2015; Song, Meng, Chen, Zhou, & Luo, 2014). As such, the ~3.5 Hz rhythm we observed when crossmodal cues mismatched with the conscious visual stimulus extends the evidence for divided attentional sampling to binocular rivalry.

We note that the issue of trial-to-trial variability when reporting on perceptual changes cannot be completely avoided in binocular rivalry research, and is important to consider. Here one might argue that variable timing in perceptual reports may blur any effects of temporal periodicity. However, our results clearly demonstrate that robust periodic cycling of perceptual states occur, despite the inherent variability in response times.

Distinct neural correlates of these attentional sampling rhythms were also found in the EEG. We found significantly greater 3.5 Hz ITPC strength for mismatched compared to matched cue types over right fronto-centro-temporal electrodes [FT8 and C6], suggesting this region may be a candidate neural correlate for divided periodic attentional sampling (Figure 3a). Accordingly, following both mismatched and matched cues, increased 3.5 Hz ITPC in this region also positively correlated with the likelihood of switching to the previously suppressed visual image across subjects (Figure 3c-d). Using visual-only stimulation, previous research has identified a pre-target ~4 Hz phase-dependency for peri-threshold perception when attention is divided across visual hemifields (Landau et al., 2015). We note that in our paradigm, attention was not divided between visual hemifields, yet interestingly, our right fronto-temporal region has previously been implicated in the reorientation of attention to unattended locations (Corbetta & Shulman, 2002; Downar, Crawley, Mikulis, & Davis, 2000; Proskovec, Heinrichs-Graham, Wiesman, McDermott, & Wilson, 2018). Taken together, our results show that periodic attentional sampling exists during binocular rivalry, when visual stimuli spatially overlap and compete for perceptual dominance.

We also found behavioral and neural correlates of focused attentional sampling during binocular rivalry when cues were consistent with the prevailing visual percept. Specifically, 8 Hz ITPC over parieto-occipital electrodes was negatively correlated with the likelihood of switching to the incongruent perceptual outcome (Figure 3e, h). Previously, phase-dependent peri-threshold perception has been reported for focused attention tasks in the visual domain (Busch & VanRullen, 2010; Busch et al., 2009; Hanslmayr et al., 2013; Mathewson et al., 2009), and has primarily implicated an approximately 7 Hz component located over fronto-central electrodes (Busch & VanRullen, 2010; Busch et al., 2009). Given the differences between paradigms, it is unsurprising that our identified region for focused attentional sampling does not coincide with those reported in previous research regarding phase-dependent perception. Nonetheless, it is notable that we identified right fronto-centro-temporal [FT8, C6] and right parieto-occipital [P6, PO8] regions in the EEG. Activity over each of these regions has previously been implicated in the reorienting of visuo-spatial attention (Corbetta & Shulman, 2002; Downar et al., 2000; Laura Dugué, Merriam, Heeger, & Carrasco, 2017; Proskovec et al., 2018), and for the integration of multisensory stimuli into a coherent percept (Beauchamp, 2005; Bushara et al., 2003; Calvert & Thesen, 2004; Driver & Noesselt, 2008; D. Zhang, Hong, Gao, Gao, & Röder, 2011). Increases in right parieto-occipital theta power (4-8 Hz) have also been shown when attending to visual stimuli in the presence of auditory distractors (van Driel, Knapen, van Es, & Cohen, 2014), with the phase of right parieto-occipital alpha (8-10 Hz) or theta (6-7 Hz) oscillations determining the perceptual outcome of bistable stimuli (Ronconi, Oosterhof, Bonmassar, & Melcher, 2017). As such, the present modulation for 8 Hz parieto-occipital ITPC is consistent with the idea that right-parietal networks may preferentially represent temporal information in the visual modality (Battelli, Pascual-Leone, & Cavanagh, 2007; Guggisberg, Dalal, Schnider, & Nagarajan, 2011). While promising, our right-lateralized response may also be due to the left-lateralized tactile input. Future experiments that control for this aspect can address this issue

Our analysis so far has revealed that when crossmodal cues mismatched the dominant binocular rivalry stimulus, that rates of attentional sampling slowed to ~ 3.5 Hz – implicating the division of attention over multiple locations. However, our exogenous cues oriented attention toward the congruency of visual and crossmodal stimuli, prompting the question: between what was attentional sampling divided? One possibility is that attentional sampling during mismatched cues was divided between two sensory modalities, as the brain tried to resolve a conflict between concurrent auditory/tactile and visual information. Figure 4a provides a schematic of this multisensory interpretation. If the neural activity in our identified region is representative of divided sampling between modalities, it constitutes the first evidence that an attentional sampling mechanism can flexibly orient between temporally co-modulating crossmodal stimuli. Although the facilitative role of attention in multisensory integration remains controversial (Hartcher-O’Brien et al., 2016; Talsma et al., 2010), we see it as a viable possibility that this mechanism resolved perceptual ambiguity through a visual perceptual switch to the competing image, rendering the multisensory stimuli congruent.

**Figure 4.**
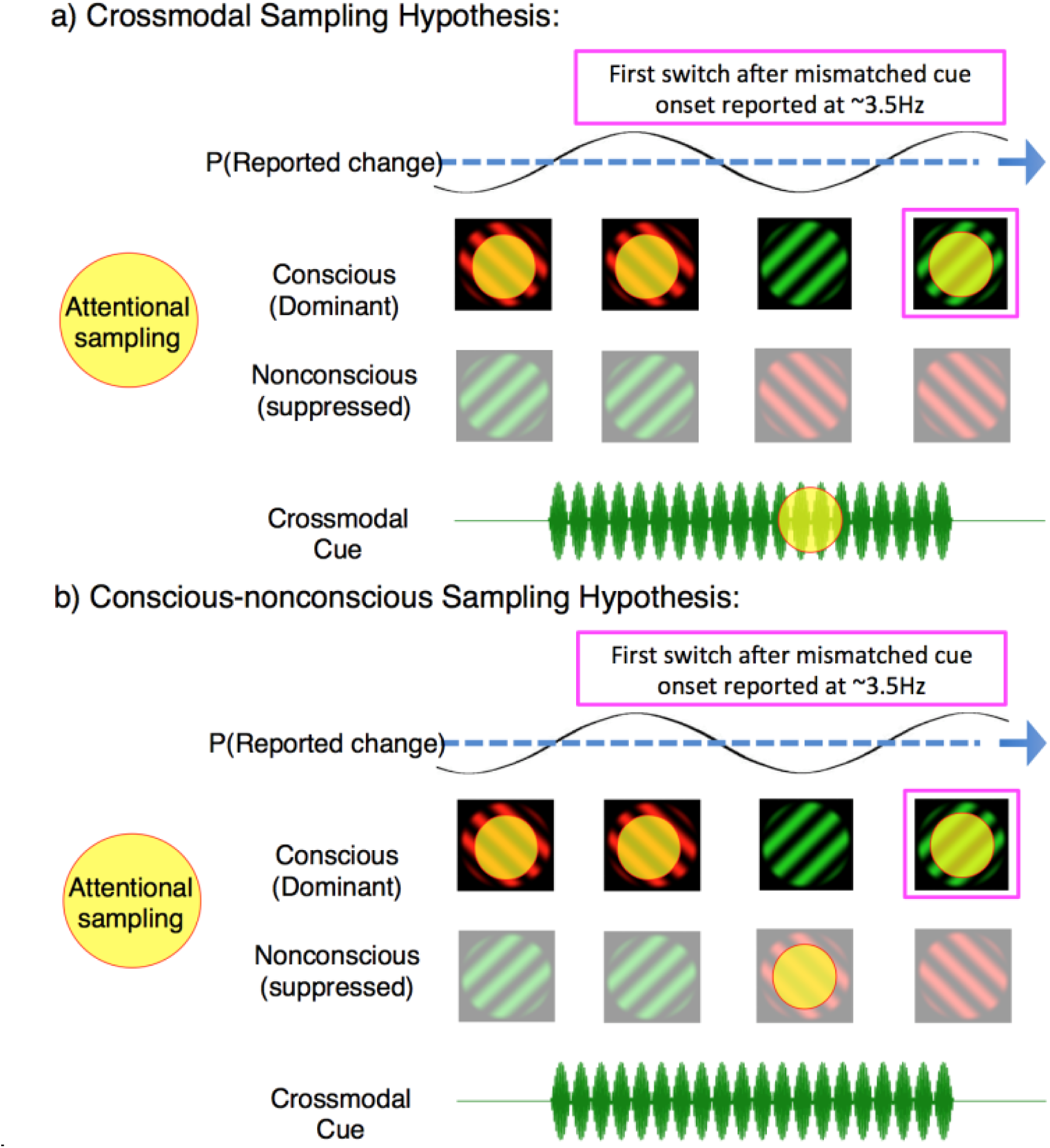
Two possible interpretations of attentional sampling during mismatched crossmodal cues. Schematic representation of attentional sampling and perceptual oscillations during binocular rivalry. a) Crossmodal sampling hypothesis: While perceiving the high-frequency visual flicker, an attended low-frequency crossmodal cue mobilises attention to sample between the dominant image and mismatched crossmodal cue at ~3.5 Hz. As a consequence, the likelihood of the first perceptual switch is modulated at ~3.5 Hz. b) Conscious-nonconscious sampling hypothesis: The onset of a mismatched cue prompts attention to sample between separate visual features, which in our paradigm consists of dominant and suppressed visual images.

Crossmodal attentional sampling may also explain why we observed low- but not high-frequency behavioral effects in the present task, particularly in the context of previous investigations regarding the binding of multisensory stimulus attributes (Fujisaki & Nishida, 2005, 2010; Lunghi et al., 2014; Vroomen & Keetels, 2010), and the limits of crossmodal temporal judgments (Fujisaki & Nishida, 2005, 2010; Holcombe, 2009; Vroomen & Keetels, 2010). For example, Fujisaki and Nishida (2005) have shown that judgments of temporal synchrony between rhythmic sensory streams degrade above ~4 Hz. It is plausible that the ineffective crossmodal cueing that we found is related to the above mentioned findings.

Having said that, one previous study using a similar design to ours was successful in eliciting a high-frequency crossmodal effect (15-20 Hz; Lunghi et al., 2014). We believe these differences are not wholly unexpected, as to optimize the present task for EEG recordings we used larger (6.5° visual angle) luminance-modulated sinusoidal gratings to facilitate subsequent steady-state visually evoked potential analyses (Davidson et al., in prep). While in comparison, Lunghi et al succeeded in showing a high-frequency effect with rivalry stimuli that were contrast-modulated narrow-band random noise patterns (3.2° visual angle), and did so under conditions analogous to our non-attend conditions. This difference in the composition of visual stimuli is noteworthy, as stimulus size is known to strongly affect rivalry dynamics (Blake, O’Shea, & Mueller, 1992). To our knowledge, whether stimulus size impacts upon crossmodal effects during binocular rivalry is unknown. However given the strength of our results for attended low-frequency flicker (Figure 1b), we note that the low- and high-frequency effects observed by Lunghi et al (2014) are not generalizable to the rivalry stimuli employed here. Similarly, whether the type of stimuli (e.g., gratings vs random noise patterns) also impacts upon crossmodal effects during rivalry represents a fruitful endeavor for research, particularly given the novel possibility of crossmodal attentional sampling.

An alternate possibility to crossmodal attentional sampling is that the 3.5 Hz rhythm in our paradigm reflects divided attentional sampling between dominant and suppressed visual images during binocular rivalry (Figure 4b). The frequency of divided attentional sampling that we observed is consistent with those obtained when visual attention has been divided between two objects or locations (Fiebelkorn et al., 2013; Landau & Fries, 2012). As our binocular rivalry stimuli necessarily occupied the same spatial location, attention in our paradigm was likely divided between either features or objects, instead of locations. Indeed, feature-based attention has already been shown to modulate neural processes when an attended target is suppressed during continuous flash suppression (Kanai, Tsuchiya, & Verstraten, 2006). During binocular rivalry, perceptual dominance is also influenced by object-based attention (Mitchell, Stoner, & Reynolds, 2004), with unconscious selection mechanisms argued to mediate perceptual transitions (Lin & He, 2009). This second alternative is also indirectly supported by the temporal limits of binocular rivalry when conflicting visual stimuli are presented asynchronously, without temporal overlap between the two eyes (O’Shea & Blake, 1986; van Boxtel, Knapen, Erkelens, & van Ee, 2008; J. van Boxtel, van Ee, & Erkelens, 2007). The maximum stimulus onset asynchrony that can sustain this type of rivalry is approximately 350 *±*50 ms, beyond which alternating stimuli introduced to one eye are perceived immediately, without rivalry occurring (van Boxtel, Alais, Erkelens, & van Ee, 2008). This limit is consistent with a 7-8 Hz attentional sampling rhythm distributed between the two conflicting stimuli (each sampled at ~3-4 Hz). When stimuli are presented rapidly enough they are temporally bound together and can engage in ongoing rivalry; when stimuli are presented slower than at 3-4 Hz, they are temporally individuated by attention, and rivalry ceases.

The suggestion that attention can sample between conscious and nonconscious vision is also consistent with a view that the underlying neuronal processes for attention and consciousness are supported by distinct neural mechanisms (Bahrami, Lavie, & Rees, 2007; Watanabe et al., 2011; for review see Tsuchiya & Koch, 2015). We note that while attentional sampling of a suppressed image suggests that attention is not sufficient for consciousness (Dehaene, Changeux, Naccache, Sackur, & Sergent, 2006; Koch & Tsuchiya, 2007; Lamme, 2003; van Boxtel, Tsuchiya, & Koch, 2010), this interpretation remains consistent with a view that attention may still be necessary for conscious perception (Chica & Bartolomeo, 2012; Cohen & Dennett, 2011; Merikle & Joordens, 1997; O’Regan & Noë, 2001; Posner, 1994, 2012).

Whether attributable to conscious-nonconscious, or visual-crossmodal attentional sampling, the present results also complement the ‘active-sensing’ hypothesis (Schroeder, Wilson, Radman, Scharfman, & Lakatos, 2010), whereby perceptual selection is determined by routine exploratory behaviors. Within active-sensing attention is critical to ‘search for’ task-relevant information from the environment (Schroeder et al., 2010), particularly via the rhythmic coordination of multisensory information (Schroeder et al., 2010; Thorne & Debener, 2014). Intriguingly, early contributions from multi-sensory (non-visual) information have been shown to determine perception (Morillon, Schroeder, & Wyart, 2015; Schroeder et al., 2010; van Atteveldt et al., 2014). While the rhythmic modulation of visual performance has also been demonstrated to follow the onset of both voluntary (Hogendoorn, 2016), and preparatory motor behaviors (Tomassini, Spinelli, Jacono, Sandini, & Morrone, 2015; Tomassini, Ambrogioni, Medendorp, & Maris, 2017). Here, in further support of the active sensing hypothesis, we have shown that task-relevant multi-sensory information can change the rhythmic modulations of perceptual selection during competition for perceptual dominance.

In summary, here we have provided novel evidence in support of attentional sampling during binocular rivalry, through the use of crossmodal cues matched to either a conscious or nonconscious visual stimulus. As the attention sampling hypothesis continues to garner traction from various psychophysical and neuronal paradigms (VanRullen, 2016a, 2016b), future targeted experimentation can confirm whether attention can indeed sample across modalities (Figure 4a), as well as if attention can sample between conscious and nonconscious neural representations during binocular rivalry (Figure 4b). The interactions between crossmodal stimuli and conscious perception represent a fruitful avenue for experimentation (Faivre, Arzi, Lunghi, & Salomon, 2017), here uncovering the previously unknown dependence of attention and consciousness on rhythmic neural dynamics of the human brain.

## Methods

### Participants

Thirty-four healthy individuals (21 female, 1 left handed, average age 23 ±4.7) were recruited via convenience sampling at Monash University, Melbourne, Australia. All had normal or corrected-to-normal vision and gave written informed consent prior to participation. Monash University Human Research and Ethics Committee approved this study, and subjects were paid 15 AUD per hour of their time, over an approximate total of 5 hours.

### Apparatus and Stimuli

Stimuli were generated using Psychtoolbox (Brainard, 1997) and custom MATLAB scripts. Each visual stimulus was viewed through a mirror stereoscope placed at an approximate viewing distance of 55 cm from computer screen (29 x 51 cm, 1080 x 1920 pixels, 60 Hz refresh rate) with the subject’s head stabilized via chin rest. Rivalry stimuli were red and green gratings displayed on a black background, with a white frame to aid binocular fusion, embedded within the wider gray background of the remaining portions of the screen. Beside each white framed image, colored arrows indicated the direction for button press (e.g., right for red, left for green). Gratings were sinusoidal with spatial frequency of 0.62 cycles per degree, oriented ± 45° from vertical, and subtended 6.5° visual angle (240 x 240 pixels on the display). Visual stimuli were contrast-modulated at either 4.5 or 20 Hz using a temporal sinusoidal envelope. The phase of each grating was static throughout each 3-minute binocular rivalry block, yet shifted after each block to reduce the effects of visual adaptation.

For crossmodal stimuli 50 Hz carrier tones were amplitude modulated by 4.5 or 20 Hz sine waves to create digital waveforms, which were either 2, 3.1 or 4 seconds in duration. For tactile stimulation, subjects clasped a wooden ball with their left hand attached to a Clark Synthesis Tactile Sound Transducer (TST429 platinum) housed in a custom sound insulated box (Lunghi et al., 2014). Auditory stimulation was delivered binaurally through Etymotic HD5 noise reduction headphones, with ACCU-Fit foam ear tips to reduce ambient noise.

### Stimulus timing

Accurate stimulus timing of synchronous visual and crossmodal stimuli was ensured with a WDM-compatible, hardware-independent, low-latency ASIO driver (www.asio4all.com), which was necessary to minimize audio buffer duration to sub-millisecond intervals and reduce latency compensation. The time-course of stimulus presentation was also physically recorded in the EEG for offline analysis. Photodiodes were used to record the flicker-envelope of visual stimuli and stored as separate channels in the ongoing EEG. The waveforms for crossmodal stimulation were simultaneously sent to both the presentation hardware and external electrode channels using a digital splitter (Redback A2630 4 Channel Headphone Distribution Amplifier). Stimulus presentation lag was assessed by computing the difference between the recorded frames of trigger-codes and actual physical trace within the EEG as part of data pre-processing. We adjusted the relative timing of behavioral and EEG data accordingly as part of this analysis. In most cases, no adjustment was necessary, requiring a maximum change of 3 frames in duration on <1% of blocks across all subjects.

### Calibration of visual stimuli

A maximum of 10 one-minute binocular rivalry blocks were performed prior to experimentation on the first day for all subjects. These blocks served to familiarize subjects with reporting their visual percepts during binocular rivalry, and to calibrate approximately equal dominance durations for the flickering stimuli in each eye. By this procedure, we instructed subjects to minimize the report of mixed rivalry and to adopt a constant criterion for perceptual report throughout the experiment (Figure S6). Contrast values for left/right eye, green/red color, and low/high frequency stimulus combinations (in total, 8 combinations) were adjusted on a logarithmic scale until approximately equivalent total dominance durations were reached (between 1:1 and 1:1.5), with the additional requirement that the average perceptual duration for each stimulus was longer than 1 second. As there were 24 unique 3-minute binocular rivalry blocks on each day of experimentation, each of the 8 combinations of visual parameters was balanced across all three crossmodal conditions.

### Calibration of auditory stimuli

Prior to experimentation, subjects were also tasked with equating the perceptual intensity of tactile and auditory stimulation for each low- and high-frequency condition, to achieve approximately equal phenomenological intensity across subjects and stimulus conditions. For all subjects, the amplitude of tactile vibrations was set to the same comfortable, supra-threshold level (approximately equivalent to 65 dB SPL). In the absence of visual stimulation, simultaneous auditory and tactile stimuli were then presented in a staircase procedure, with subjects adjusting the amplitude of auditory tones to match the perceived intensity of simultaneous tactile vibrations. They performed the matching task separately within low-frequency auditory tones and tactile vibrations and within high-frequency auditory tones and tactile vibrations. This calibration procedure was performed on each day of testing, to account for differences in the insertion depth of inner-ear headphones across separate days.

### Experimental Procedure and Behavioral Analysis

Twentyfour three-minute binocular rivalry blocks were presented on each of the two separate days of testing. In each block, subjects reported their dominant visual percept during rivalry while receiving occasional crossmodal cues, which were either auditory, tactile, or simultaneous auditory and tactile. In a given three-minute block, we presented only one of the three types of crossmodal cues. The order of these blocks were randomized for each subject and each day of experimentation. In each block, 12 trials of crossmodal cues were presented. Each cue was either low (4.5 Hz) or high (20 Hz) frequency auditory and/or tactile stimulation. Six cues were presented for each frequency, with durations composed of three x 2 s, two x 3.1 s, and one x 4 s cues. To increase uncertainty of the timing of the cues, we inserted three null cues (which we call visual-only periods, Figure 1) without any crossmodal stimulation for a duration of 2.6 s (the average of crossmodal cue durations). We also used these visual-only periods as baseline for behavioral analysis (Figure 1 and 2). We randomized the order of all cues, which were separated with uniform jittering by 7-10 s ISI within each block.

Across all sessions, subjects were told to focus on accurately reporting their dominant visual percept at all times via button press. As the state of the button-press was sampled at 60 Hz, the same rate as the video refresh rate, we were able to estimate the probability and time-course of binocular rivalry dynamics over 16.7 ms intervals. Over two sessions on separate days, subjects distributed attention between visual rivalry and crossmodal cues based on separate task instructions. On Day 1 for *n*= 18 or Day 2 for *n*=16, subjects were instructed to ignore the crossmodal cues and to focus on reporting only visual rivalry. For the other session, subjects were instructed to distribute attention across both visual rivalry and crossmodal cues. To ensure their attention was on task, these alternate days included task instructions for subjects to silently tally the number of times the temporal frequency of an attended crossmodal cue matched that of their dominant visual percept at the time of crossmodal cue’s offset. Due to the varied duration of crossmodal cues, this task ensured that attention was allocated consistently throughout the presentation of crossmodal cues. To familiarize subjects with these task demands, an additional two practice blocks (three minutes each) were included during the calibration procedure on the relevant day of experimentation. Although 34 subjects were retained for final analysis, five others were recruited and began the experiment, yet failed to complete their second day of experimentation. One other subject was removed due to their failure in following task instructions and excessive movement during EEG recording.

### Evaluation of attention-on-task

To evaluate the attentional allocation to both visual and crossmodal stimuli, at the end of each 3-minute block we asked subjects to verbally report their subjective estimate of the number of crossmodal stimuli which were matched in temporal frequency to the flicker of their dominant visual percept at the point of attended-crossmodal cue offset. Then, we defined an index, ‘attention to cues’ (Figure 1c, x-axis) as the correlation coefficient between 24 subjective estimates (one per attended block) and the actual recorded occurrences of congruent stimuli. Supplementary Figure S7 displays the correlation between subjective and actual congruent stimuli for an exemplary subject.

### Behavioral data analysis

We preprocessed the button press data to accurately estimate the timing of changes in visual consciousness during binocular rivalry. First, we categorised each time-point according to the flicker frequency of the dominant visual stimulus reported. To analyze the time-course of the probability of a button press state (Figure 1b), we categorized button-presses (which could be either low or high-flicker) as either congruent or incongruent with the ongoing crossmodal stimulus frequency. Then, we obtained the probability of a congruent button press state as a function of time per subject, by averaging responses at each time point across all 144 trials per attention x frequency cue subtype. For visual-only periods, the left button (corresponding to left-eye dominance) was arbitrarily set to congruent prior to the averaging of probability traces within subjects. As visual parameters were balanced across all blocks, this selection necessarily balanced across visual frequency and colour parameters, and we note that the identical analysis performed using right-eye congruence produced equivalent results. Mismatched (Figure 1d) or matched (Figure 1e) condition comparisons were then defined by whether the congruent button (left-eye dominant) was pressed at cue onset. In these two panels, we set the y-axis for ‘Probability of seeing mismatched flicker’, to reflect the probability of perceptual states that differ in temporal frequency from the crossmodal cue. In Figure 1b, d, and e, we compared among conditions (visual-only, low-frequency or high-frequency crossmodal cue x attended vs unattended conditions) with one-way repeated-measures ANOVAs. We defined significant differences among conditions at those time points that survived corrections for multiple comparisons with planned comparisons between cue types and the visual-only baseline, using FDR at q = .05 (Benjamini & Yekutieli, 2001).

### Perceptual Switch Index (PSI)

To quantify crossmodal effects during binocular rivalry, we defined the perceptual switch index (PSI). PSI is the difference in the probability of a change in percept when comparing attended low-frequency to all other crossmodal cues. For the y-axis in Figure 1c, we calculated the PSI as the difference in the probability of viewing a congruent visual flicker over the period 1-4 s after stimulus onset. The same subtraction was used to compare the probability of viewing the previously suppressed visual flicker following mismatched (Figure 3c, g) and matched cues (Figure 3d, h), for the period 2-4 s after onset. This shorter time window was selected to capture when the crossmodal effects on binocular rivalry emerged for mismatched and matched cues. A similar pattern to the results displayed in Figure 3 was shown when a wider window was used (e.g. 0-4 s, data not shown).

### Spectral analysis of first switches

For our spectral analysis (Figure 2), we focused on the first perceptual switches, which were the first time-point recording a change in button-press state after cue onset. To account for individual variation in the amount of overall switches, the proportion of switches at each time point was first calculated at the subject level, before averaging across all subjects. We sampled button presses at 60 Hz (or every 16.7 ms). For the spectral analysis of perceptual switches (Figure 2f), we applied a single-taper fast Fourier transform (FFT) to the period 0.5 - 2 seconds after cue onset (Nyquist = 30 Hz, a half bandwidth = 0.67 Hz). This window was selected to restrict the analysis so that all the analyzed trials occurred during an attended cueing period (as the minimum crossmodal cue duration was 2 seconds), and to remove transient button presses occurring early in the cue period, which were unlikely to be caused by crossmodal match or mismatch (Figure S2). We display the frequency range of 0 - 15 Hz for all conditions, as no higher frequencies (15-30 Hz) were significant after our two-stage statistical criteria.

### Statistics on spectra of first switch timing

To assess the statistical significance of behavioral spectra we used a two-stage statistical testing procedure as applied in previous investigations of attentional sampling (Landau & Fries, 2012) and electrophysiological research (Maris & Oostenveld, 2007). At the first stage, we first detected significant frequencies (at *p* < .005 uncorrected) through a non-parametric randomization procedure. After obtaining the spectral amplitude for the observed data across subjects, we generated a null distribution of first switches during the same cue period by randomly shifting switch-times within each subject, while keeping the number of perceptual switches the same. We generated 5000 surrogate datasets in this way, to test the null hypothesis that there were no temporal effects on the timing of perceptual switches. We then compared the amplitude of the Fourier transform from the observed and the surrogate data at each frequency. We regarded the spectral amplitude at a certain frequency to be significantly above chance, if the observed spectral amplitude exceeded the top 99.5% of the null-distribution of amplitudes at each frequency generated by surrogate data.

At the second stage, we applied a cluster criterion, which corrects for multiple comparisons across multiple frequencies through a permutation procedure (Maris & Oostenveld 2007). We required that the first-level significance (*p* < .005 uncorrected) be sustained for at least two neighboring frequencies, and retained the sum of their clustered test-statistics (amplitudes in this case) as our observed data. Then, from our surrogate dataset, we calculated the maximum cluster-based amplitudes per surrogate (maximum spectral amplitude excluding 0-1 Hz and nearest neighbor), which we retained as the null-distribution to compare against our observed data. Candidate clusters survived this second order analysis when their observed cluster-based test-statistics exceeded the top 95% of the null distribution, or corrected to *p_cluster_* < .05 if testing across multiple clusters. The null-distributions obtained for our frequencies of interest in Figure 2f are shown in Figure S3.

### EEG recording and analysis

EEG was recorded at a sampling rate of 1000 Hz using three BrainAmp amplifiers and 64-channel ActiCap (BrainProducts), with the impedance of each electrode kept below 10 kΩ Ground and reference electrodes were AFz and FCz at recording, respectively. After re-referencing the data to the average of all channels, we performed linear detrending and bandpass filtering (0.1- 60 Hz with a Hamming-windowed finite impulse response filter) and down-sampled the data to 250 Hz before time-frequency analysis.

We performed all time-frequency analyses using the Chronux toolbox (http://chronux.org; Bokil, Andrews, Kulkarni, Mehta, & Mitra, 2010) and custom MATLAB scripts. To resolve our frequencies of interest (especially between 3.5 and 4.5 Hz), we used a single-taper Fourier transform with a time-window of 2 seconds, which resulted in a half bandwidth (*W*) of 0.5 Hz (*W* = 1/2). This half bandwidth is consequently capable of resolving differences between 3.5 and 4.5 Hz, as demonstrated in Figure 3b and f.

### ITPC analysis

To assess the neural correlates of attentional sampling (Figure 3), we analyzed the inter-trial phase coherence (ITPC) within electrodes, over multiple time-frequency points(Bastos & Schoffelen, 2016). ITPC is an amplitude-normalized measure of the degree to which EEG responses are phase locked to the onset of an exogenous cue, ranging between 0 (random phase over trials) and 1 (perfect phase coherence over trials). To compute ITPC, the consistency of phase angles is computed as the length of the average of unit phase vectors in the complex plane over trials. For a given time, t, and frequency, f,

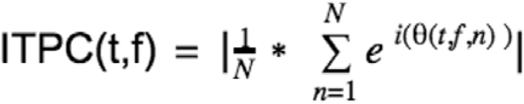

where N is the number of trials, and θ is the phase angle at time t, frequency f, and trial n.

Due to the stochastic nature of perceptual alternations during binocular rivalry, the number of available trials for analysis in each mismatched and matched cue type ranged from 12 to 36 trials across subjects (after averaging first across subjects, the mean number of trials was 24 (±1.5) trials across matched / mismatched and attention conditions). Because the bias level (or expected chance level for pure noise data) of ITPC is strongly influenced by the number of trials, we took additional measures to equate the number of mismatched and matched cue types for the analysis. To achieve this aim, the maximum number of trials recorded for a given cue combination was identified across subjects. Following this, subjects with fewer numbers of trials had their observed number of trials supplemented by resampling with replacement from their recorded trials, equating them to the predefined maximum for each condition. Upon this resampled dataset, the ITPC was computed, and this process repeated 100 times. Our analysis confirmed that upsampling, compared to downsampling, reduced the bias introduced when equating ITPC values across subjects. As the difference in ITPC between auditory, tactile, and combined auditory and tactile cues was not significant, we proceeded by combining crossmodal cue types within each subject.

### ITPC statistics

To investigate the neural correlates of attentional sampling, we analysed evoked ITPC, the increase in ITPC during 0 to 2 s after onset compared to −2 to 0 s before onset. Similar to our statistical approach for the behavioral spectral analysis described above, we used a two-stage statistical testing procedure for this analysis. At the first stage, we performed a *t*-test (two-tailed) comparing whether evoked 3.5 and 8 Hz ITPC differed between mismatched and matched conditions across subjects at each electrode. At each electrode, we used the mean evoked ITPC value obtained from the upsampling method described above. As a result of the *t*-tests, if we found a cluster of at least two neighboring electrodes with *p* < .05 (uncorrected), where inter-electrode distance did not exceed 3.5 cm, we proceeded using this cluster in the second stage of statistics. As a result of this cluster criterion, we always identified a minimum size of 2-electrode clusters (Figure 3a and e).

At the second stage, we first computed the sum of observed *t*-scores within the identified cluster, which we retained as our observed test-statistic (Figure S5). To create the null distribution, condition labels (mismatch and match) were randomly shuffled for each electrode within each subject, to create two surrogate datasets the same size as our original mismatch and match conditions. Then the *t*-scores were computed for each electrode based on our surrogate datasets, and the electrode with the maximum *t*-score and the maximum *t*-score of its neighbors retained. The sum of these *t*-scores were then retained per shuffle, and this procedure repeated 2000 times to obtain a null distribution of the sum of *t*-scores around the maximum electrode for each shuffle of our surrogate data. Against this distribution, the sum of observed *t*-scores for the candidate cluster was then compared. When the observed sum of *t*-scores was within the top 5% (or cluster corrected to *p* < .05) then we concluded that there was a significant difference between mismatch and match conditions. The null-distributions and observed test-statistics produced by this analysis are shown in Figure S5.

### Data Availability

The data in this study are available from the corresponding author upon request, and will be available on Figshare upon online publication.

### Author Contributions

D.A., N.T., and J.J.A.V.B. conceived of the initial extension of Lunghi et al (2014) into an EEG paradigm. D.A. contributed tactile-stimulation equipment and code for auditory stimulus construction. M.J.D designed and programmed the research experiment reported in this article, with feedback from N.T. and J.J.A.V.B. M.J.D collected the experimental data. M.J.D performed the initial behavioral and EEG analysis, with feedback from J.J.A.V.B. and N.T. M.J.D conceived of and performed the attentional sampling analysis with feedback from N.T. and J.J.A.V.B. M.J.D. wrote the article, with feedback from D.A, N.T. and J.J.A.V.B.

## Acknowledgements

M.J.D. was supported by an Australian Government Research Training Program Scholarship. D.A. was funded by an Australian Research Council Discovery Project (DP150101731). N.T. was funded by an Australian Research Council Future Fellowship (FT120100619) and Discovery Project (DP130100194). The authors thank Bryan Paton and Claudia Lunghi for technical advice, and Brandon Lam for early piloting of the experimental paradigm.

---

**Supplemental Figure 1, related to Figure 2.**
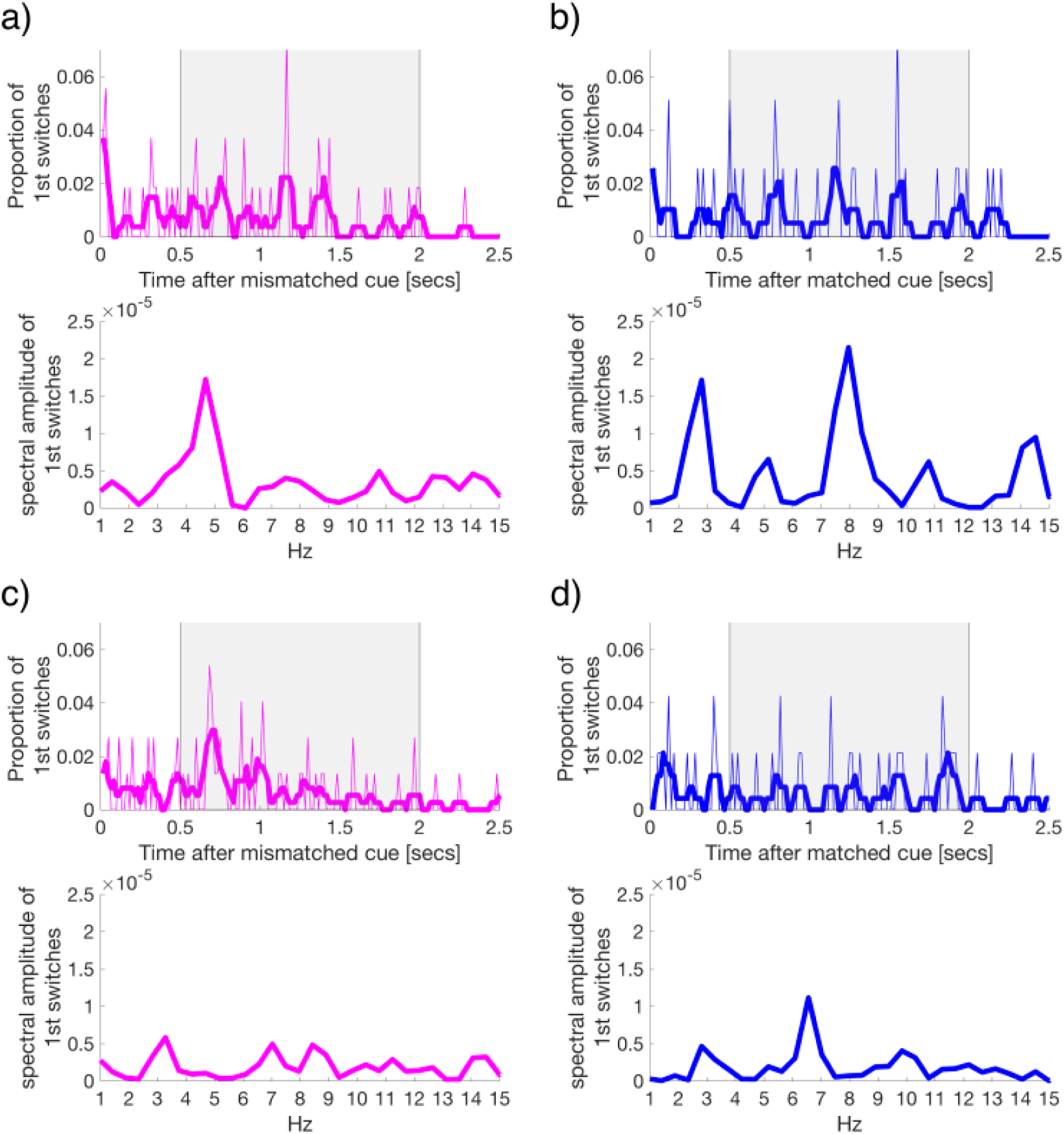
Representative subject data demonstrating that the strength of attentional sampling varies across subjects. Perceptual switches recorded during binocular rivalry after attended low-frequency cues either a,c) mismatched, or b,d) matched with the dominant percept at cue onset. a-d)Top panels show the first perceptual switches after crossmodal cue at each time-point, binned over 16.7 ms intervals. Bottom panels show the spectra of these time-courses following FFT within the grey-shaded time window, 500-2000 ms after cue onset.

**Supplemental Figure 2, related to Figure 2.**
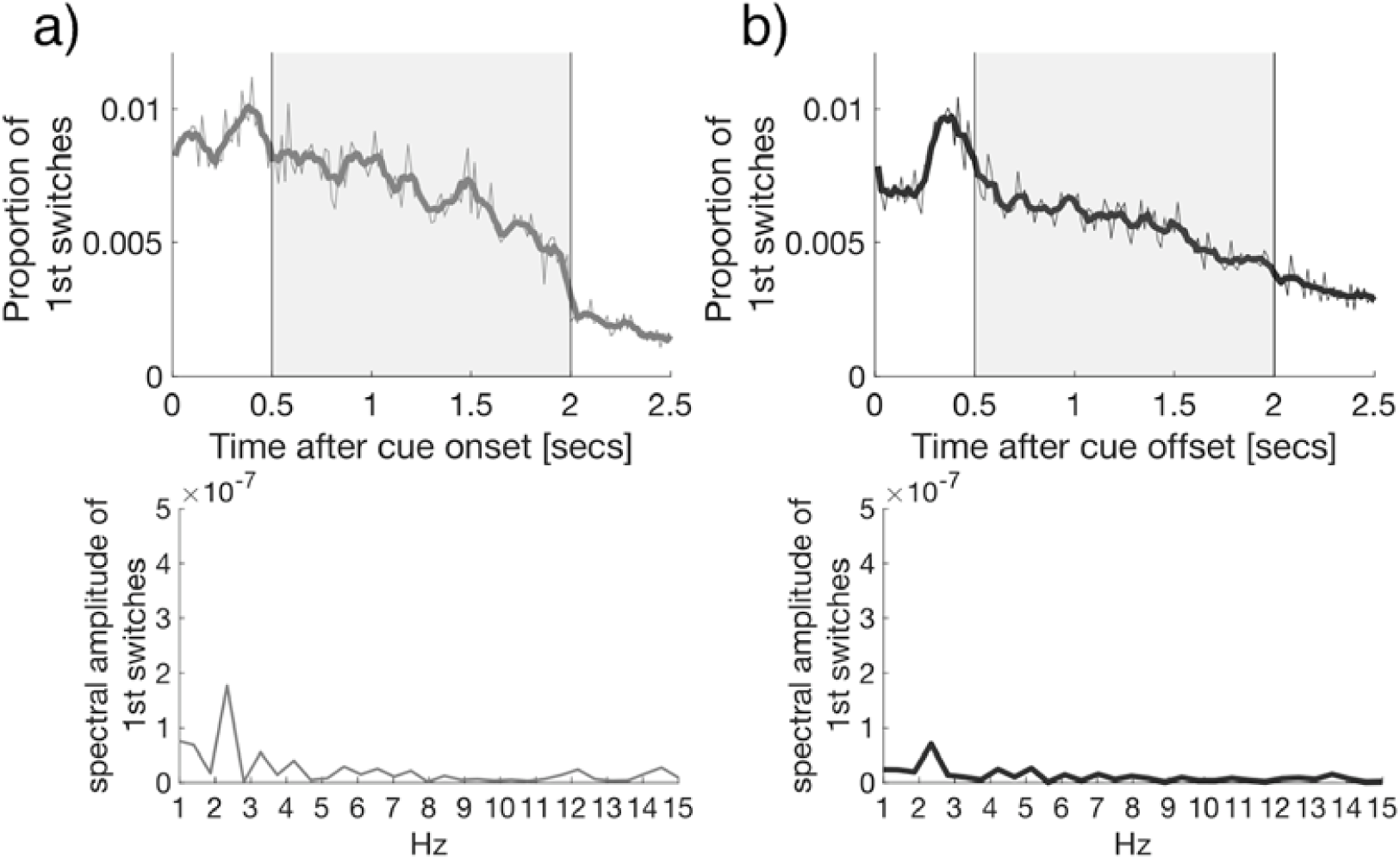
The time course of the proportion of first switches made at each time point; following any crossmodal cue onset (a), and following crossmodal cue offset (b). An analysis of the time-course of perceptual switches reveals no significant spectral peaks at 3.5 or 8 Hz. Y-axis scaled as per Figure 2f. The presence of an early peak (0 - 0.5 s) in the proportion of first switches suggests that these changes may be due to stimulus transients, rather than the cue-conditional allocation of attention. As such this early time-window was omitted from subsequent analysis.

**Supplemental Figure 3, related to Figure 2f.**
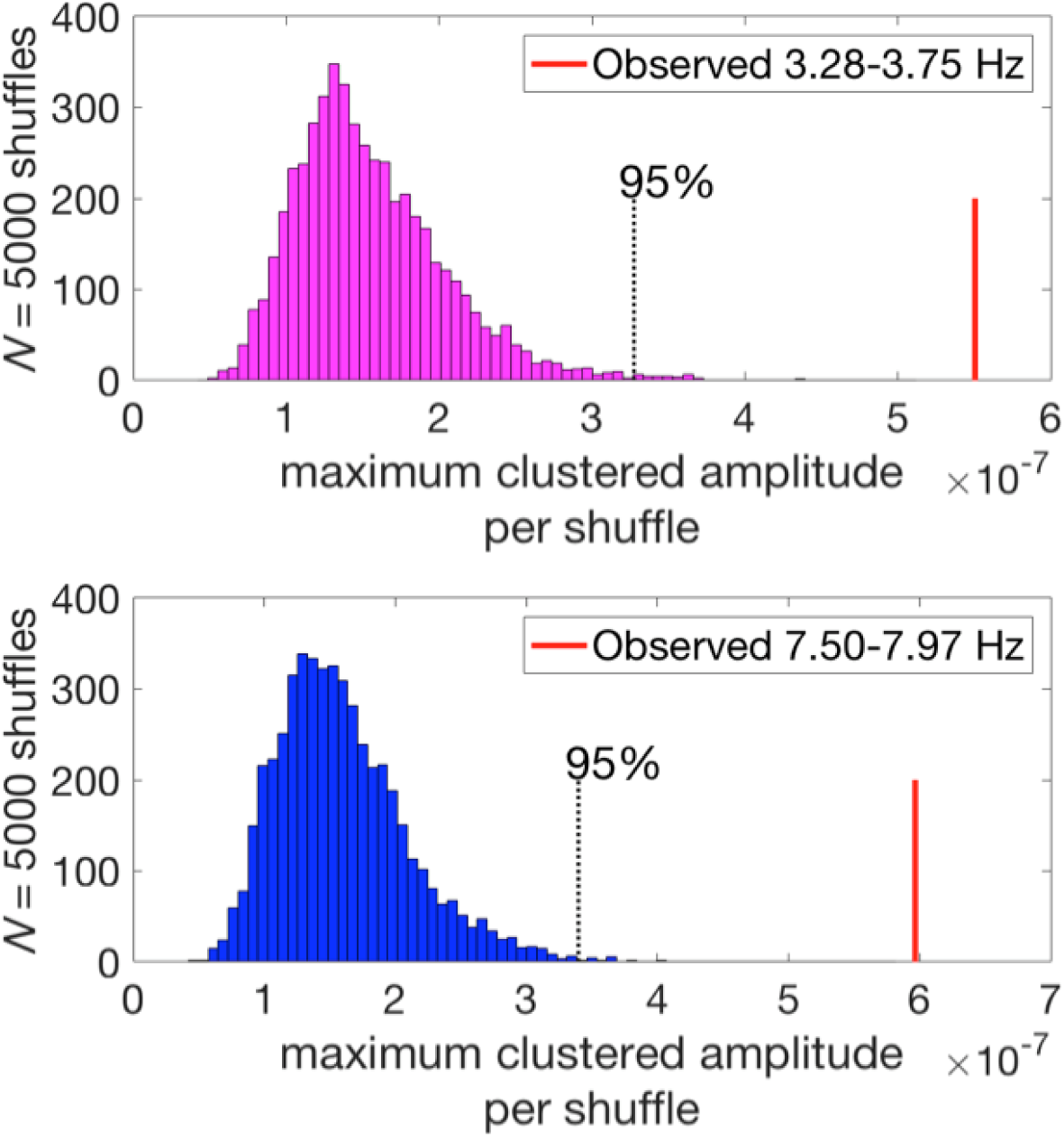
The null-distributions for the surrogate datasets generated by the randomization procedure, and the actually observed values of second-stage statistics (i.e., maximum and its highest neighbor’s summed Fourier amplitude). After satisfying first-level criteria (p < .005 uncorrected for two neighboring frequencies), we proceeded to this second-stage statistical test. The observed second-stage statistics (red line) were regarded as significant after cluster corrections for multiple comparisons at p < .05 level; exceeding the top 95% of the null distribution.

**Supplemental Figure 4, related to Figure 2.**
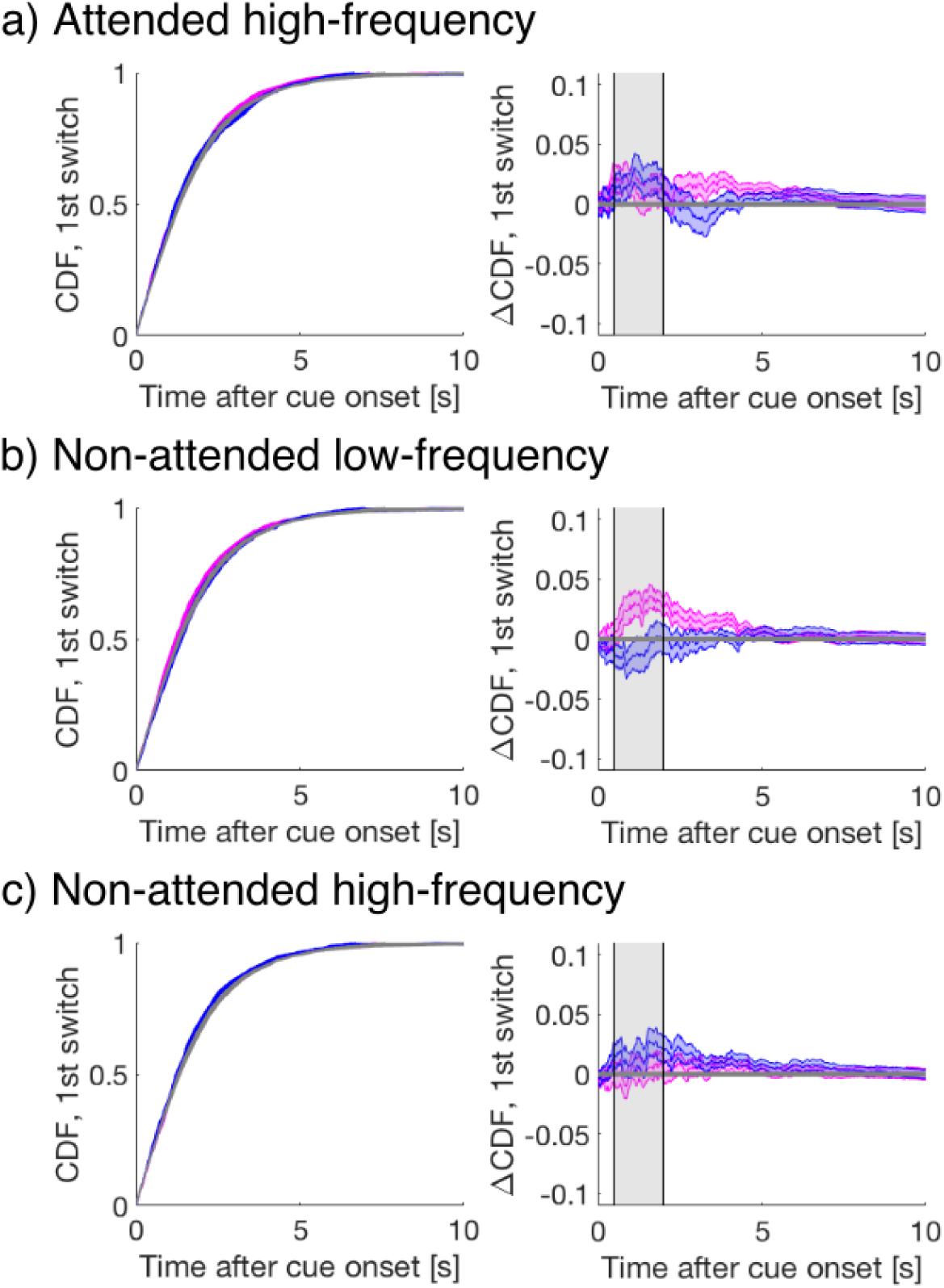
Left column) The cumulative density function (CDF) of the time to first-switch for all conditions other than attended low-frequency. Mismatched, matched, and visual-only conditions are colored in magenta, blue, and grey in all panels. Lines and shading show mean and standard error across subjects (N=34). Right column) the difference in CDF between conditions, each of which failed to exhibit any significant crossmodal effects on perceptual switches compared to visual only periods (FDR q = .05). Thus, we did not pursue further spectral or neural analyses of these conditions.

**Supplemental Figure 5, related to Figure 3.**
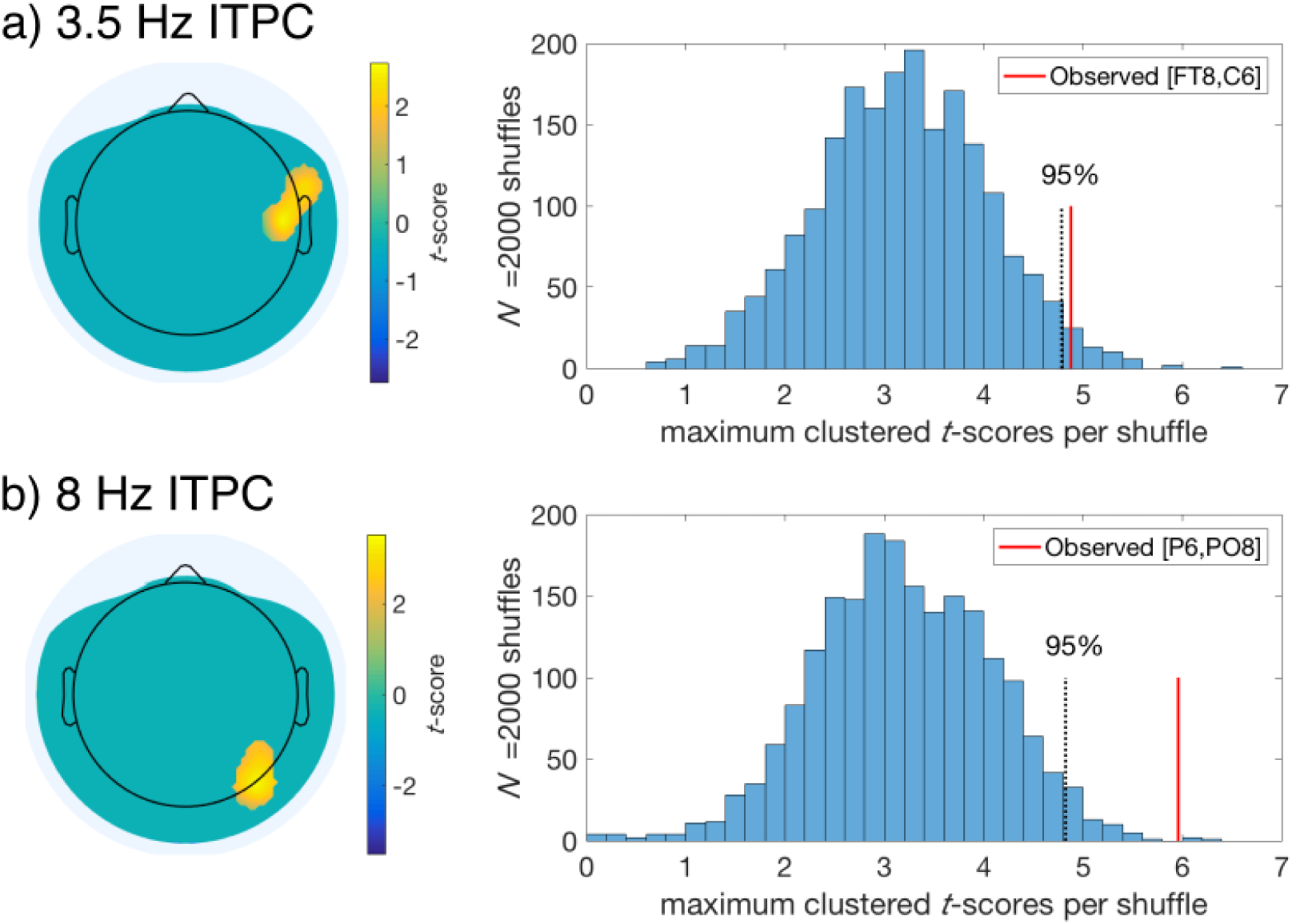
Displayed are the regions selected for correlation analysis after satisfying our two-stage statistical tests on evoked ITPC, comparing mismatched and matched conditions for a) 3.5 Hz and b) 8 Hz. The right panels display the resulting null distributions obtained after the permutation of condition labels (mismatched vs matched) and performing t-tests across subjects on the mean evoked ITPC for each electrode after upsampling (see Method). The maximum clustered t-scores per shuffle were retained to create the null distributions. The observed sum of t-scores is displayed as a vertical red line, while the top 95% of the distribution is marked with a vertical dotted back line.

**Supplemental Figure 6.**
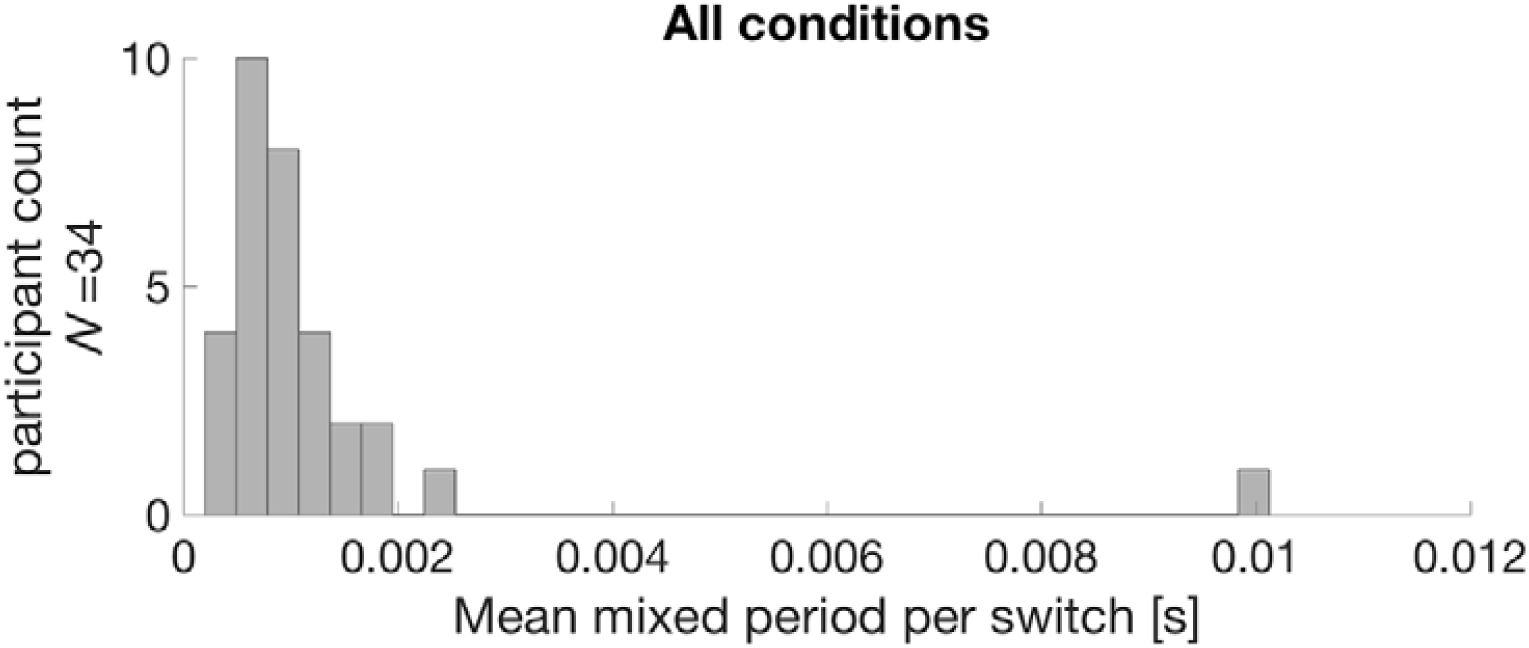
Across all experimental periods, the average duration of mixed periods per switch per subject was less than 16.7 ms (our binning width), thus showing that mixed percepts are unlikely to have contributed to an increase in the variance of perceptual report timing. Switches happened instantly, with zero or one mixed frame (16.7 ms) on average.

**Supplemental Figure 7, related to Figure 1c.**
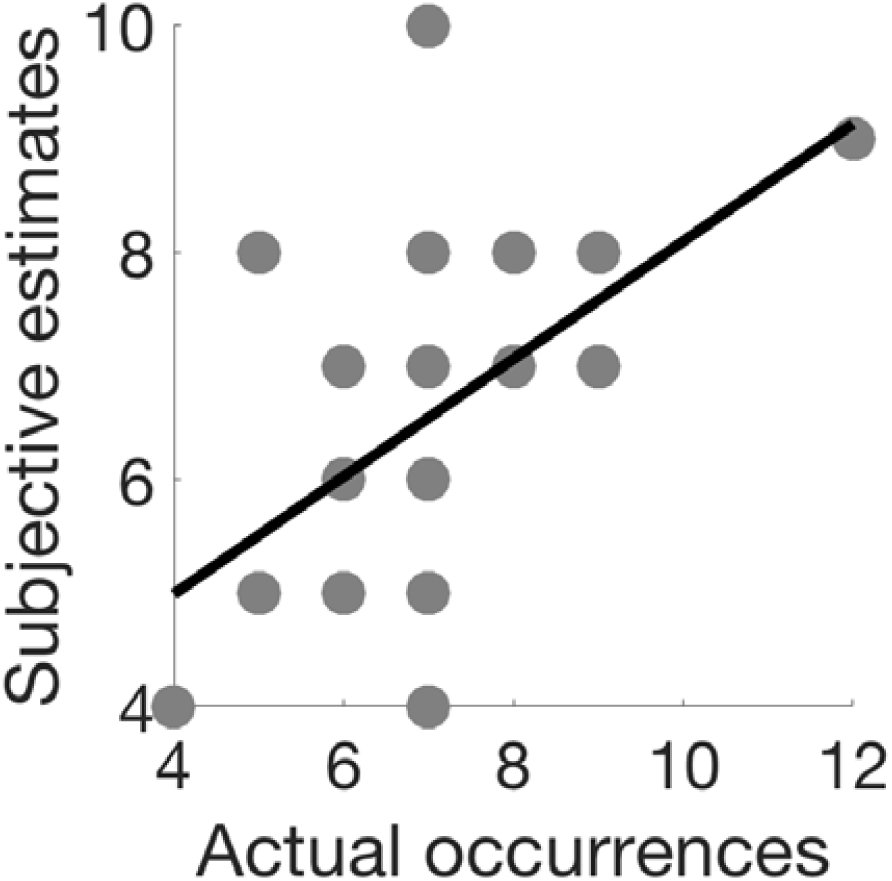
Definition of “attention to cues” in Figure 1c. Y- and x-axes of this figure are the subjective and actual congruent crossmodal and visual stimuli in the attended sessions from one exemplary subject. We defined the correlation coefficient between the two (here, r = .55) as the ‘attention to cue’ index used as x-axis in Figure 1c.

